# A comparison between power spectral density and network metrics: an EEG study

**DOI:** 10.1101/614271

**Authors:** Matteo Demuru, Simone Maurizio La Cava, Sara Maria Pani, Matteo Fraschini

## Abstract

Power spectral density (PSD) and network analysis performed on functional correlation (FC) patterns represent two common approaches used to characterize Electroencephalographic (EEG) data. Despite the two approaches are widely used, their possible association may need more attention. To investigate this question, we performed a comparison between PSD and some widely used nodal network metrics (namely strength, clustering coefficient and betweenness centrality), using two different publicly available resting-state EEG datasets, both at scalp and source levels, employing four different FC methods (PLV, PLI, AEC and AEC_C_). Here we show that the two approaches may provide similar information and that their correlation depends on the method used to estimate FC. In particular, our results show a strong correlation between PSD and nodal network metrics derived from FC methods (PLV and AEC) that do not limit the effects of volume conduction/signal leakage. The correlations are less relevant for more conservative FC methods (AEC_C_). These findings suggest that the results derived from the two different approaches may be not independent and should not be treated as distinct analyses. We conclude that it may represent good practice to report the findings from the two approaches in conjunction to have a more comprehensive view of the results.

## 1. Introduction

Power spectral density (PSD) and network analysis performed on functional correlation (FC) patterns represent two common approaches used to characterize Electroencephalographic (EEG) and Magnetoencephalographic (MEG) time-series data [1]. Despite these two approaches are widely used, the possible association between the results derived from their application is probably overlooked and may deserve more attention. Although few attempts to characterize this association have been performed, investigating, for instance, the dependency between patterns of global synchrony (phase) and local synchrony (amplitude) [2] on synthetic data, usually the two approaches are not analyzed in conjunction. In this study we aim to understand to what extent the two approaches differ in experimental data and if the corresponding analyses may be interpreted as completely separated and independent. In order to investigate in more details their possible relationship, we performed a comparison between PSD analysis and some widely used nodal network metrics (namely strength, clustering coefficient and betweenness centrality), using two different publicly available resting-state EEG datasets. To assess potential limitations due to scalp-level analysis [3,4], the analysis was further replicated using a source level approach. In order to control the possible effects derived from the use of different FC methods, which may result from distinct neural mechanisms [5], we performed the analysis using four different techniques to estimate patterns of phase- and amplitude-based correlation: the Phase locking value (PLV) [6], the Phase lag index (PLI) [7], the Amplitude Envelope Correlation (AEC) [8] and a corrected version performing a time-domain orthogonalization procedure (AEC_C_) [8]. Our analysis was focused on the alpha band since it has been previously shown to provide the largest signal to noise ratio and the more reliable estimate of FC networks [9].

## 2. Material and methods

### 2.1 EEG datasets

Two different EEG datasets were used for the analysis. The first dataset (EEG_DS1) is the EEG motor movement/imaginary dataset [10,11] (https://www.physionet.org/pn4/eegmmidb/), a freely available set of 64 channels EEG recordings, consisting of several tasks including one-minute eyes-closed resting-state from 109 subjects. The second dataset (EEG_DS2) is another freely available set of 64 channels EEG recordings [12,13], consisting of eyes-closed resting-state from 12 subjects. All the analysis was performed using five epochs of 12 seconds [14] for each subject extracted from one-minute of eyes-closed resting state condition. All the reported results refer to the investigation of the alpha frequency band (8 – 13 Hz). In order to evaluate the consistency of our results between scalp and source analysis [3], we replicated the analysis at source-level using source-reconstructed time-series obtained by using Brainstorm software (version 3.4) [15], using the protocol described by Lai and colleagues [3].

### 2.2 Features extraction

For each subject and each epoch, we have extracted a set of features from the EEG time-series. In particular, the relative alpha band power was computed for each channel (at scalp level) and for each ROI (at source level) as the ratio between the sum of the original PSD (computed using the Welch method in Matlab R2017b) over the frequency range in 8 – 13 Hz and the sum of the original PSD over the frequency range in 1 – 40 Hz (total power). Later, four different and common methods to estimate (alpha band) FC patterns were used: PLV [6], AEC [8], PLI [7] and AEC_C_ [8] (an orthogonalized version of AEC). Finally, using the BCT [16], a set nodal network metrics were computed from the FC patterns: strength (the sum of weights of links connected to the node), clustering coefficient (the fraction of triangles around a node) and betweenness centrality (the fraction of all shortest paths in the network that contain a given node). All the extracted nodal features were represented as feature vectors of 64 (at scalp level) or 68 (at source level) entries. All the code used to perform the analysis is available at the following link: https://github.com/matteogithub/PSD_NET_comparison.

### 2.3 Statistical analysis

In order to estimate the relevance of the correlations between the extracted PSD-based and network-based features, the Spearman’s rank correlation coefficient (rho) was computed for each comparison at channel level, without performing average across subjects, epochs or channels.

## 3. Results and discussion

The results show a clear association between the power analysis, namely the alpha relative power content, and the network analysis performed using the four different FC methods and computing the three different nodal metrics. The level of association varies depending on the specific FC method and on the computed nodal metrics, but it is persistent over the three tested scenarios: EEG_DS1, EEG_DS2 at scalp level and EEG_DS1 at source level. In particular, the association is more evident for the strength and the clustering coefficient, whereas it results lower for the betweenness centrality. As shown in Figure 1, for the EEG_DS1, the association results in the range 0.698 – 0.224 for the strength, with a pick rho value for the PLV (rho = 0.698). The PLV based network metrics confirm a higher association with alpha relative power also for the clustering coefficient (rho = 0.736) and for the betweenness centrality (rho = - 0.344).

**Figure 1.**
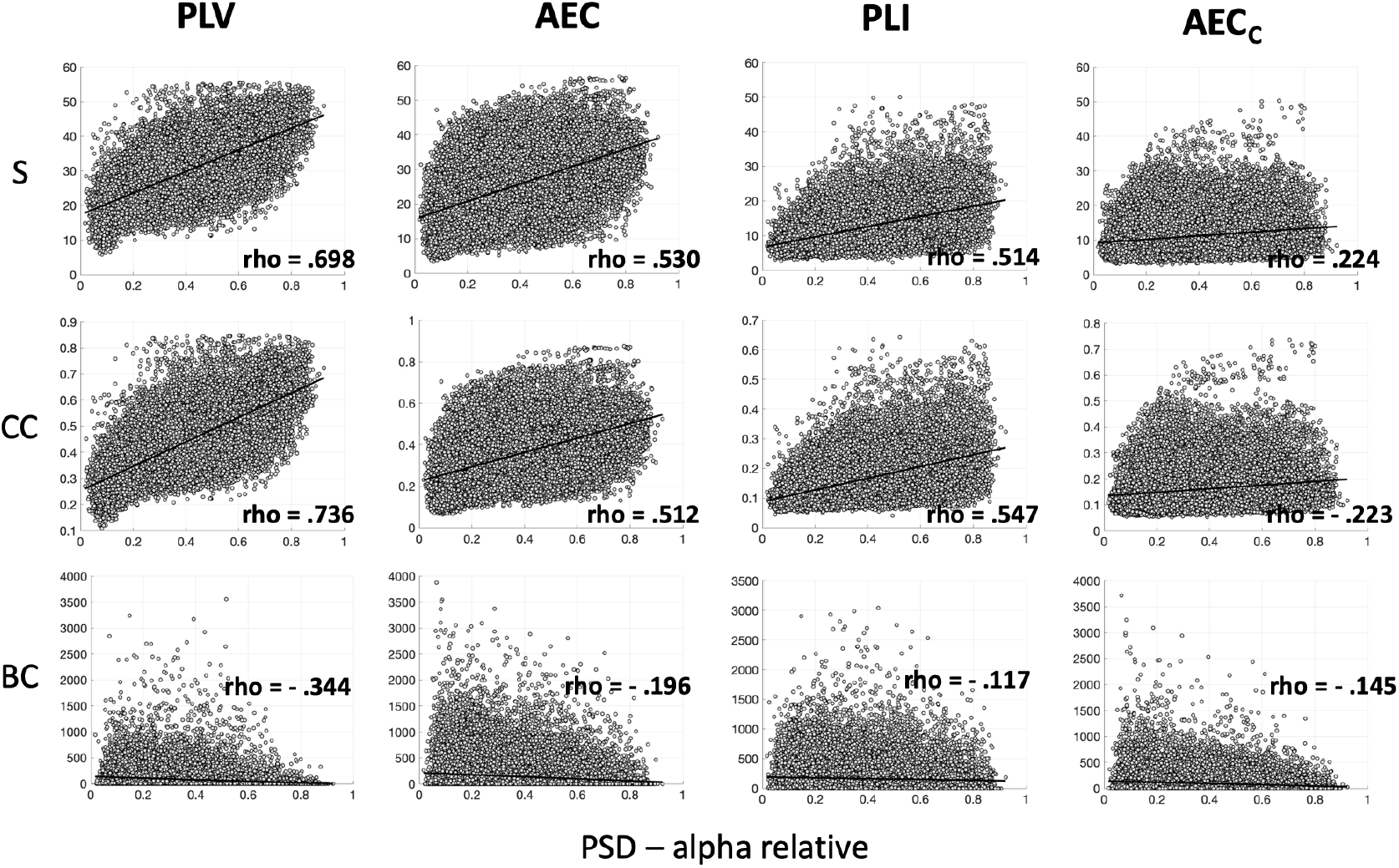
Scatter plots and correlations between alpha relative power and nodal network metrics for all the FC methods using the EEG_DS1 at scalp level. S, CC and BC respectively represent strength, clustering coefficient and betweenness centrality.

Considering the relative high association between the two approaches, we have replicated the whole analysis using a different EEG dataset (EEG_DS2) comparable with the previous one (EEG_DS1) in terms of number of channels and experimental condition. Even the replication, although with different correlation values, show a clear association between the two approaches. As shown in Figure 2, not only the results still show relative high correlation values, with highest magnitude of rho equals to 0.493 for the strength and 0.572 for the clustering coefficient, but even more importantly, the reported findings are consistent with the previous analysis in term of the effects due to the different FC methods. The lowest level of association was obtained when using the AEC_C_ methods to extract the FC patterns.

**Figure 2.**
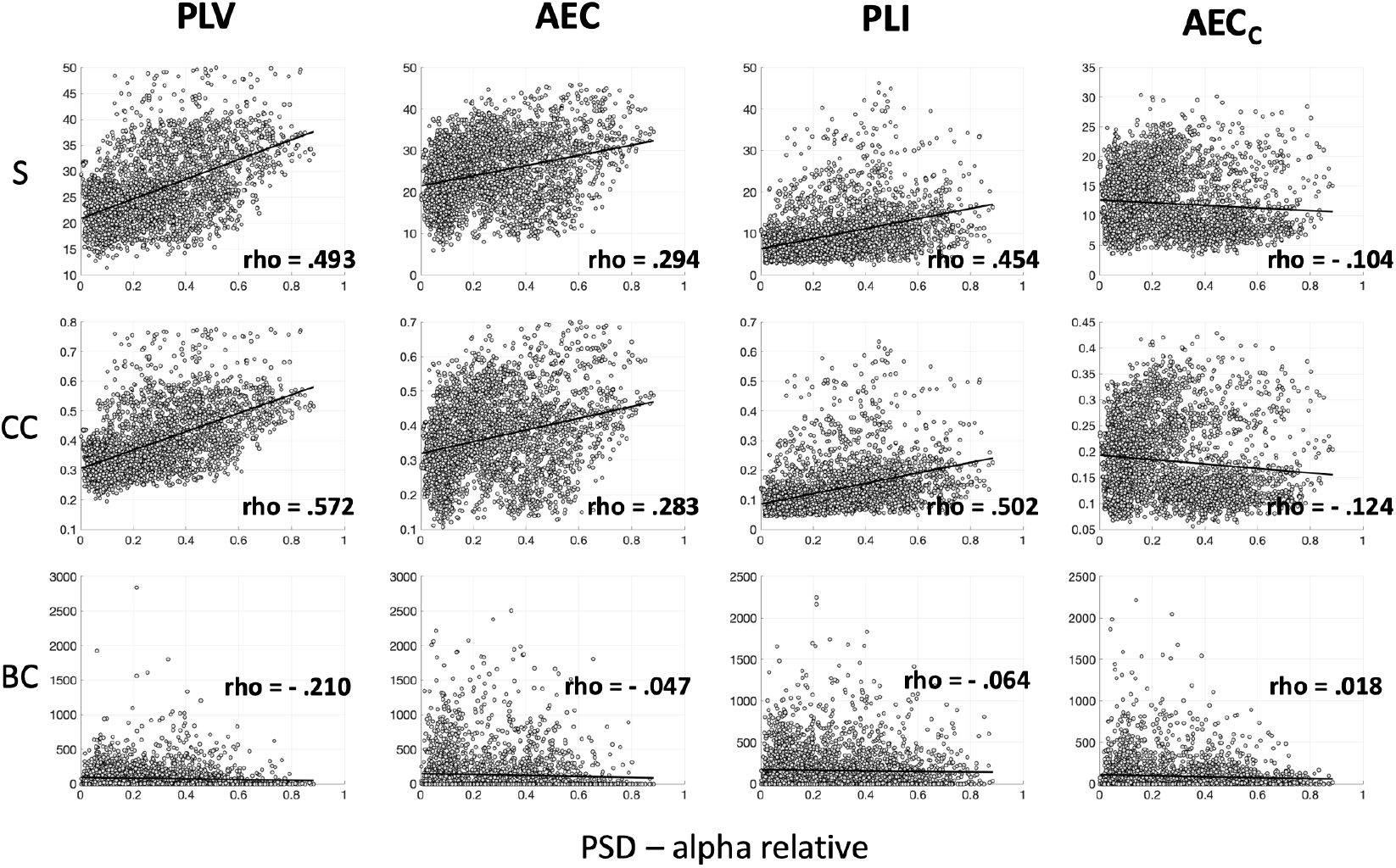
Scatter plots and correlations between alpha relative power and nodal network metrics for all the FC methods using the EEG_DS2 at scalp level. S, CC and BC respectively represent strength, clustering coefficient and betweenness centrality.

Finally, considering the problems related with the not straightforward interpretation of results derived from an EEG scalp level analysis, we have further replicated the analysis with source-reconstructed time-series (derived from EEG_DS1) using the procedure as described in [3]. Even in this case, as shown in Figure 3, the results are still in line with the previous analysis performed at scalp level. In this latter case, where the source-based FC patterns should be clearly less affected by volume conduction and signal leakage, the differences among the FC methods are even less evident. For both the strength and the clustering coefficient, the associations remain moderately high, respectively in the range 0.618 – 0.328 and 0.601 – 0.316. Despite for PLV the correlation with PSD is not surprising, as already reported in previous studies [17,18], the correlation between the power and the other FC methods, as for PLI [19], is less straightforward. In summary, these findings show that the results obtained using the two approaches, namely power analysis and network analysis derived from the use of FC methods, may be correlated and therefore should not be treated as completely independent. In particular, assessing and reporting the relationship between the two analyses may avoid to overestimate and to overinterpret their results when treated as separated approaches. More in general, despite network tools are easily accessible, widely used and provide appealing interpretations, findings derived from their application should be interpreted with caution and not translated as a direct consequence of functional brain mechanisms and/or alterations, as shown with simple simulations [20].

**Figure 3.**
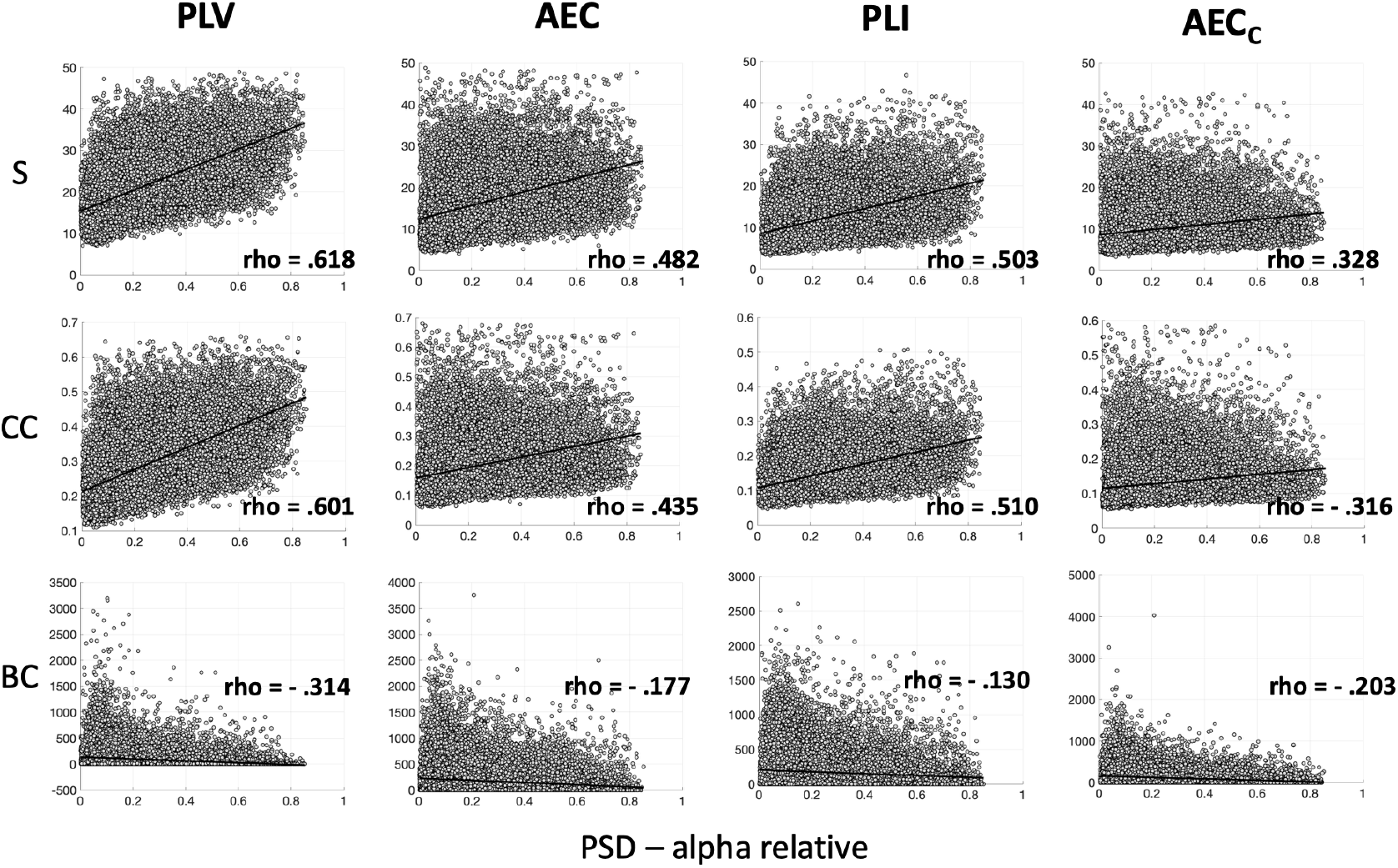
Scatter plots and correlations between alpha relative power and nodal network metrics for all the FC methods using the EEG_DS1 at source level. S, CC and BC respectively represent strength, clustering coefficient and betweenness centrality.

## 4. Conclusions

In conclusion, this study suggests that the results derived from the two different approaches, PSD and network analysis, may be strongly associated. The level of association may depend on the specific FC method used to estimate the patterns of interactions and it is evident at both scalp and source level. As a consequence, we think that it would represent a good and necessary practice to report the results from the spectral analysis in conjunction with those obtained from network analysis.

## Acknowledgements

Simone Maurizio La Cava was supported by Regione Autonoma della Sardegna POR FESR Sardegna 2014-2020 Asse 1 Azione 1.2.2 CC F26C18000500006. Matteo Fraschini was in part supported by Regione Autonoma della Sardegna research project “Algorithms and Models for Imaging Science [AMIS]”, FSC 2014-2020 - Patto per lo Sviluppo della Regione Sardegna.

